## Supplemental Figures and Tables for "On the Emergence of P-Loop NTPase and Rossmann Enzymes from a Beta-Alpha-Beta Ancestral Fragment"

**Supplemntary Information**

**for**

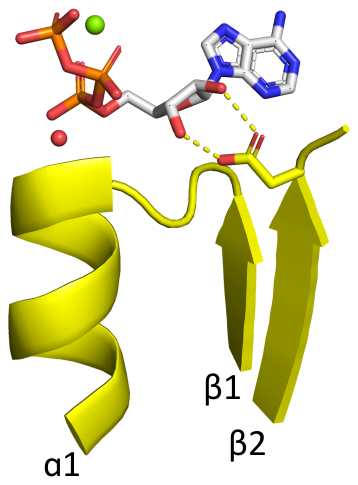

**Figure S1. Rossmann domain binding ATP in the canonical binding mode.** The phosphate binding loop of ECOD domain e3h5nA5, a Rossmann domain from the ECOD F-group 2003.1.9.15, binds the nucleotide ATP in a largely canonical fashion, including the bidentate interaction between the β2-Asp and the ribose hydroxyls (see **Main Text**). The conserved water characteristic of the Rossmann fold is shown as a red sphere. An Mg^2+^ cation is shown as a green sphere. Other F-groups with canonical binding of an NTP include 2003.1.9.10 and 2003.1.4.3.

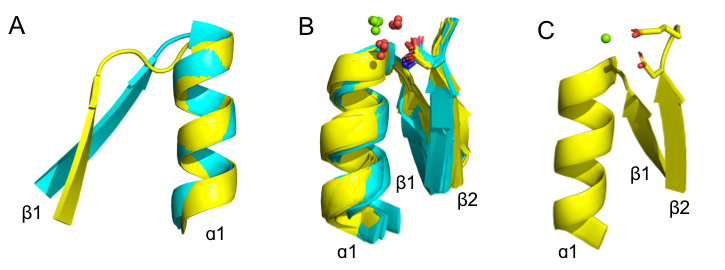

**Figure S2. Features of the tubulin binding site (see also Table S1).** **A**. The tubulin phosphate binding loop resides between β1 and α1 (cyan; ECOD domain e5j2tA1) as in all Rossmanns, yet is shorter and more compact than the canonical Rossmann binding loop (yellow; ECOD domain e1lssA1). **B**. The conserved β2 Asp (yellow structures; ECOD domains e1ffxB2, e1sa1C2, e2btoA2, e2hxfA2, e3cb2A2, e3e22C2, e3r4vA1, e3zbqA2, e4ffbA4, e4ffbB1) can also be replaced by asparagine in some structures (cyan structures; ECOD domains e1rq2A2, e1w5fA2, e2r6r11, e2vamA1, e2vapA2, e2xkbC5, e3v3tA3, e3zidA1, e4b45A3, e4b46A1, e4dxdA1, e4e6eA1, e4m8iA2). This residue is positioned to interact with the catalytic magnesium (green spheres), typically via a water molecule (red spheres).

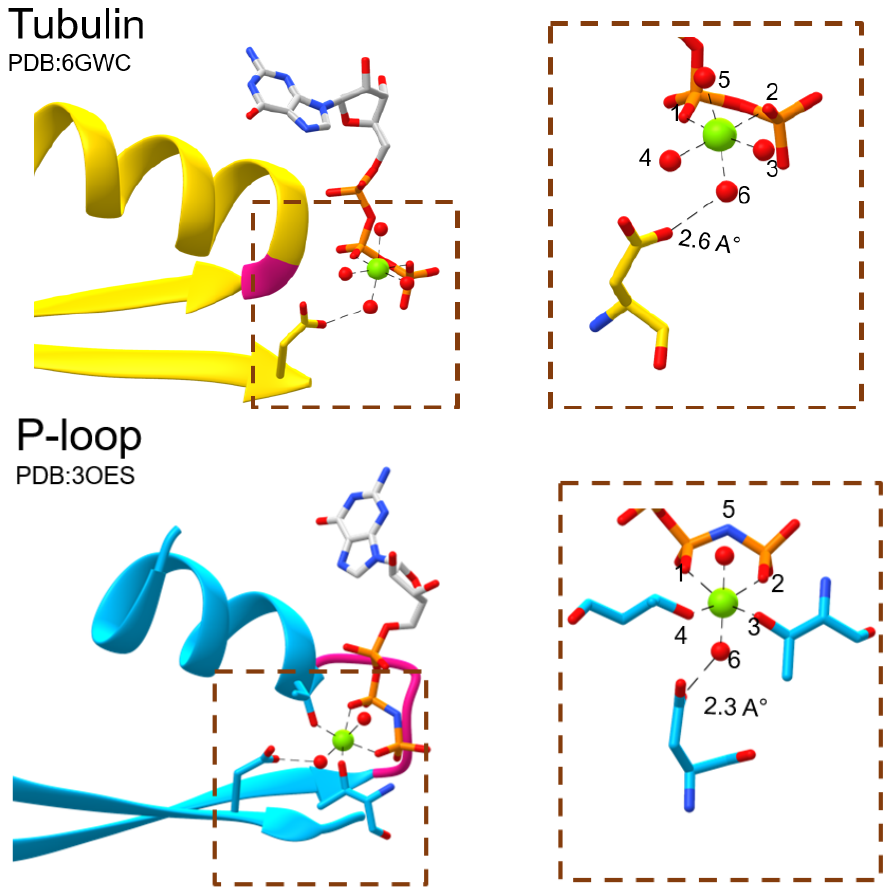

**Figure S3. The Tubulin β2-Asp and the P-Loop Walker B interact with waters that occupy equivalent sites around the catalytic Mg^2+^ cation.** The catalytic Mg^2+^ cation in both Tubulin (top panel) and P-loop domains (bottom panel) forms octahedral coordination complexes. Both the β2-Asp of Tubulin and the Walker B of P-Loop interact with the water at position 6 of the coordination sphere. Water molecules rendered as red spheres and Mg^2+^ cations rendered as green spheres.

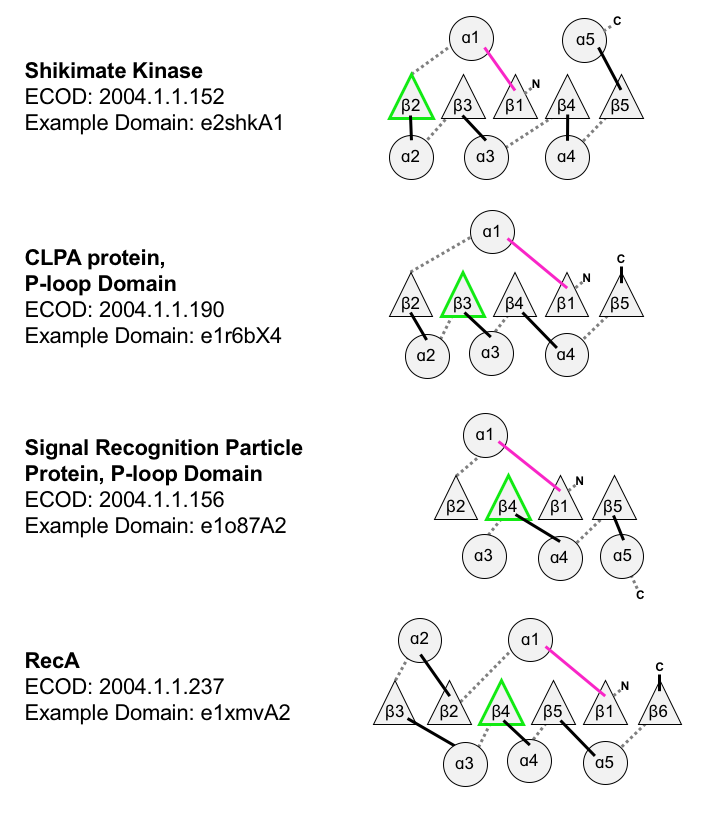

**Figure S4**. **Topological diversity in the P-loop evolutionary lineage.** In addition to the examples noted here, which all have parallel β-sheets, the P-loop lineage also has instances of anti-parallel strands inserted into the β-sheet (see **Figure 4C**, main text). The loop bearing the Walker A and Walker B motifs are colored magenta and green, respectively.

**Table S1: Dication binding in tubulins**

| **PDB ID** | **Chain ID** | **Tubulin Subfamily** | **Bound Metal** | **β2 Residue** | **Distance to Bound Metal^*^** |
| --- | --- | --- | --- | --- | --- |
| 4F6R | A | Alpha-beta tubulin | Mg | Asp | 3.9 |
| 4FFB | A | Alpha-beta tubulin | Mg | Asp | 4.2 |
| 4I4T | A | Alpha-beta tubulin | Mg | Asp | 4 |
| 4U3J | A | Alpha-beta tubulin | Mg | Asp | 4 |
| 5IYZ | A | Alpha-beta tubulin | Mg | Asp | 4.1 |
| 5NQU | A | Alpha-beta tubulin | Mg | Asp | 4 |
| 6GWC | A | Alpha-beta tubulin | Mg | Asp | 4.1 |
| 1W58 | A | FtsZ | Mg | Asn | 4.7 |
| 1W5A | A | FtsZ | Mg | Asn | 4.5 |
| 1W5F | A | FtsZ | Mg | Asn | 4.5 |
| 2R75 | A | FtsZ | Mg | Asn | 4.3 |
| 1Z5V | A | Gamma tubulin | Mg | Asp | 3.7 |
| 1Z5W | A | Gamma tubulin | Mg | Asp | 4 |
| 3RB8 | A | PhuZ | Mg | Asp | 3.9 |
| 2XKA | A | TubZ | Mg | Asn | 5.3 |
| 2XKB | A | TubZ | Mg | Asn | 5 |

^*^Taken as the distance between the metal and the closest sidechain oxygen of the β2-Asp or the sidechain oxygen for the equivalent Asn.

**Table S2. Summary of Rossmann proteins that share a bridging theme with the P-Loop domain e1ko7A1.**

|  | Best alignment  of N-segment* | | Aligned Theme | | Best alignment of C- segment* | |  |
| --- | --- | --- | --- | --- | --- | --- | --- |
| ECOD Domain | Score (bits) | #residues | Score  (bits) | #residues | Score  (bits) | #residues | ECOD  F-Group |
| e1uayA1 | -2.666667 | 2 | 42 | 68 | 17 | 63 | 2003.1.1.417 |
| e4qecA1 | -3.666667 | 3 | 52.333333 | 113 | -16 | 13 | 2003.1.1.418 |
| e2p68B1 | 4 | 14 | 64 | 118 | -11 | 10 | 2003.1.1.419 |
| e1hdcA1 | -2.666667 | 1 | 30 | 47 | 33.666667 | 82 | 2003.1.1.420 |
| e3rkuB1 | -6 | 15 | 39 | 46 | 40 | 83 | 2003.1.1.421 |
| e1xu9A1 | -6.333333 | 11 | 56.333333 | 111 | -7.666667 | 20 | 2003.1.1.422 |
| e4iboD1 | 26.666667 | 75 | 15.666667 | 34 | 13.666667 | 38 | 2003.1.1.423 |
| e1ulsB1 | -2.666667 | 4 | 43 | 49 | 38.666667 | 83 | 2003.1.1.424 |
| e4nbwA1 | -1.333333 | 3 | 30.333333 | 75 | 18 | 53 | 2003.1.1.425 |
| e3gedA1 | -2.666667 | 2 | 42.333333 | 78 | 14.333333 | 52 | 2003.1.1.426 |
| e3v8bA1 | -2.333333 | 4 | 38.666667 | 49 | 34.333333 | 85 | 2003.1.1.427 |
| e2bgkA1 | 31.333333 | 66 | 23.666667 | 34 | 17.333333 | 42 | 2003.1.1.428 |
| e3awdA1 | 32.333333 | 122 | 15.333333 | 21 | -26 | 4 | 2003.1.1.429 |
| e2p91A1 | 19.666667 | 110 | 27.333333 | 35 | -15 | 8 | 2003.1.1.430 |
| e4trrG1 | -3.333333 | 1 | 34 | 32 | 47.666667 | 109 | 2003.1.1.431 |
| e4bmvC1 | -0.333333 | 5 | 28.666667 | 25 | 64.333333 | 113 | 2003.1.1.432 |
| e4is3C1 | 22.333333 | 64 | 12.666667 | 23 | 21 | 53 | 2003.1.1.433 |
| e3gk3A1 | 0 | 72 | 21 | 60 | -2.666667 | 16 | 2003.1.1.434 |
| e1iy8A1 | 2.666667 | 6 | 28.666667 | 31 | 51.666667 | 107 | 2003.1.1.435 |
| e4zd6C1 | 34.333333 | 103 | 21.666667 | 31 | -30 | 2 | 2003.1.1.436 |
| e1g0nB1 | -2 | 2 | 29.333333 | 30 | 51 | 109 | 2003.1.1.437 |
| e1zmoA1 | 8 | 98 | 17.333333 | 26 | -1.333333 | 25 | 2003.1.1.438 |
| e2c07A1 | 27.333333 | 100 | 17.333333 | 28 | 8.333333 | 25 | 2003.1.1.439 |
| e3n74B1 | 4 | 107 | 11 | 21 | 2.666667 | 21 | 2003.1.1.440 |
| e1hxhA1 | 25.333333 | 67 | 23 | 38 | 17.333333 | 43 | 2003.1.1.441 |
| e4kwhA1 | 18.333333 | 55 | 6.333333 | 32 | 27 | 53 | 2003.1.1.442 |
| e2ehdA1 | -0.666667 | 2 | 58 | 99 | 13.333333 | 34 | 2003.1.1.332 |
| e3asuB1 | -5 | 1 | 39.333333 | 50 | 30 | 77 | 2003.1.1.333 |
| e3ftpA1 | 3.333333 | 8 | 40.666667 | 72 | 27.333333 | 53 | 2003.1.1.334 |
| e4allA1 | 23.666667 | 104 | 36 | 43 | -21.666667 | 4 | 2003.1.1.335 |
| e4z0tA1 | -3.333333 | 0 | 42 | 95 | 15.333333 | 37 | 2003.1.1.336 |
| e3u9lA1 | -2.666667 | 3 | 41.333333 | 77 | 10.666667 | 54 | 2003.1.1.337 |
| e1n5dA1 | 24.333333 | 92 | 29.666667 | 33 | -7.666667 | 24 | 2003.1.1.338 |
| e4j1sA2 | 43.666667 | 65 | 33.666667 | 42 | 3.666667 | 41 | 2003.1.1.339 |
| e4di7A1 | 32 | 83 | 23.333333 | 30 | -49.333333 | 1 | 2003.1.1.340 |
| e1qsgA1 | 13.666667 | 130 | 19.666667 | 21 | -29 | 2 | 2003.1.1.341 |
| e3tjrB1 | 4.333333 | 7 | 32 | 31 | 47.333333 | 110 | 2003.1.1.342 |
| e3guyH1 | 23.333333 | 73 | 23.666667 | 49 | 2.666667 | 21 | 2003.1.1.343 |
| e4h15A1 | 21.666667 | 68 | 23 | 60 | -5.333333 | 21 | 2003.1.1.344 |
| e2pd3A1 | 17.333333 | 111 | 18.333333 | 35 | -19 | 8 | 2003.1.1.345 |
| e4m87A1 | 26 | 110 | 24 | 35 | -16.666667 | 8 | 2003.1.1.346 |
| e2vz9B11 | 16.333333 | 60 | 11.666667 | 31 | 16 | 53 | 2003.1.1.347 |
| e1ooeA1 | 15.666667 | 113 | 12 | 29 | -15.666667 | 7 | 2003.1.1.348 |
| e3oifA1 | 28.333333 | 120 | 8.666667 | 29 | -22.333333 | 4 | 2003.1.1.349 |
| e1ek6A1 | -1.666667 | 5 | 20.333333 | 47 | 30 | 77 | 2003.1.1.410 |
| e3enkA1 | 0.333333 | 6 | 52.333333 | 68 | 22.333333 | 70 | 2003.1.1.411 |
| e1z45A2 | -2 | 3 | 36.666667 | 73 | 4.666667 | 55 | 2003.1.1.412 |
| e4lisA1 | -0.333333 | 3 | 30.333333 | 55 | 9.666667 | 76 | 2003.1.1.413 |
| e1kvtA1 | -3.333333 | 2 | 36 | 53 | 31.666667 | 82 | 2003.1.1.414 |
| e3ondA1 | 0.333333 | 9 | 39 | 108 | 0.333333 | 13 | 2003.1.1.11 |
| e1li4A1 | 29 | 94 | 18.333333 | 27 | 1.333333 | 19 | 2003.1.1.12 |

* N-segment relates to the segment preceding the identified bridging theme, and C-segment to the segment following it.
